# Do try this at home: Age prediction from sleep and meditation with large-scale low-cost mobile EEG

**DOI:** 10.1101/2023.04.29.538328

**Authors:** Hubert Banville, Maurice Abou Jaoude, Sean U.N. Wood, Chris Aimone, Sebastian C. Holst, Alexandre Gramfort, Denis-Alexander Engemann

## Abstract

EEG is an established method for quantifying large-scale neuronal dynamics which enables diverse real-world biomedical applications including brain-computer interfaces, epilepsy monitoring and sleep staging. Advances in sensor technology have freed EEG from traditional laboratory settings, making low-cost ambulatory or at-home assessments of brain function possible. While ecologically valid brain assessments are becoming more practical, the impact of their reduced spatial resolution and susceptibility to noise remain to be investigated. This study set out to explore the potential of at-home EEG assessments for biomarker discovery using the brain age framework and four-channel consumer EEG data. We analyzed recordings from more than 5200 human subjects (18-81 years) during meditation and sleep, focusing on the age prediction task. With cross-validated *R*^2^ scores between 0.3 - 0.5, prediction performance was within the range of results obtained by recent benchmarks focused on laboratory-grade EEG. While age prediction was successful from both meditation and sleep recordings, the latter led to higher performance. Analysis by sleep stage uncovered that N2-N3 stages contained most of the signal. When combined, EEG features extracted from all sleep stages gave the best performance, suggesting that the entire night of sleep contains valuable age-related information. Furthermore, model comparisons suggested that information was spread out across electrodes and frequencies, supporting the use of multivariate modeling approaches. Thanks to our unique dataset of longitudinal repeat sessions spanning 153 to 529 days from eight subjects, we finally evaluated the variability of EEG-based age predictions, showing that they reflect both trait- and state-like information. Overall, our results demonstrate that state-of-the-art machine learning approaches based on age prediction can be readily applied to real-world EEG recordings obtained during at-home sleep and meditation practice.

## 1 Introduction

In recent years, the emergence of low-cost mobile electroencephalography (EEG) devices has made it possible to monitor and record brain activity in entirely new environments, dramatically improving access to the technology for various applications (Johnson and Picard, 2020; Krigolson et al., 2017; Niso et al., 2022). Wearable EEG devices typically have few channels, are wireless and use dry electrodes but are significantly more affordable than their clinical counterparts. This makes them a perfect candidate for translating EEG screening procedures to at-home settings or wherever clinical or research infrastructure is not available. The accessibility and affordability of low-cost mobile EEG devices can open the door to the tracking of brain activity and health on a day-to-day basis. Still, a major obstacle to this goal is the need for large volumes of labeled clinical EEG data to develop pathology-specific predictive models that can deliver actionable biomarkers, *i*.*e*., measurable characteristics of the brain that provide information about its function and pathology (Ward, 2010).

To address this challenge, it is worthwhile to explore applying pretext tasks to learn representations (Banville et al., 2021) or proxy measures (Dadi et al., 2021) in the context of wearable EEG. These techniques can distill biomedical information of interest in the scarcity or absence of direct labeled data and are related to transfer learning (Pan and Yang, 2010). In human neuroscience, predicting age from brain data has emerged as a popular blend of these concepts with the potential to deliver biomarkers of brain health and pathology by capturing deviations from what “normal” brains “look like” (Cole and Franke, 2017; Cole et al., 2018; Denissen et al., 2022). In this framework, the *brain age delta* ∆ is defined as the difference between an age estimate (*i*.*e*., the age predicted by a model trained on a healthy population) and chronological age. A positive ∆ can reflect premature aging or pathology that make affected brains “look” older by comparison, *e*.*g*., caused by Alzheimer’s disease or disturbed sleep (Chu et al., 2023; Cole and Franke, 2017; Cole et al., 2018; Franke and Gaser, 2012; Lee et al., 2022; Weihs et al., 2021). Evidence on the clinical utility of brain age keeps accumulating. Most of the research involved has focused on structural magnetic resonance imaging (MRI), thus restricting brain age measurement to clinical and research settings, potentially neglecting opportunities to explore this technique in at-home and ambulatory settings outside MRI scanners, and severely limiting the feasibility of obtaining subject-wise repeated measures.

A recent line of research has provided first hints that the information obtained through functional neuroimaging modalities could provide similar, or even complementary, information on brain age (Engemann et al., 2020; Liem et al., 2017; Xifra-Porxas et al., 2021). Specifically, EEG-based brain age modeling with EEG has been explored during resting state (Al Zoubi et al., 2018; Vandenbosch et al., 2019), clinical screening (Sabbagh et al., 2020, 2022) and sleep (Brink-Kjaer et al., 2020; Paixao et al., 2020; Sun et al., 2019; Ye et al., 2020). In light of these findings, low-cost mobile EEG emerges as a promising tool for estimating brain age out-of-the-lab as part of a regular at-home screening procedure. For example, the Muse S headband (Hashemi et al., 2016) facilitates the study of brain activity in personal environments during meditation exercises and sleep.

In this work, we analyzed consumer-grade EEG recordings from more than 5200 individuals to explore the feasibility of a real-world brain age metric. As clinical or cognitive-testing information was absent in the present study, our work focuses on establishing a methodology and benchmarks for brain age prediction from at-home EEG recordings. Specifically, we addressed the following questions: 1) Can age be predicted from low-cost mobile EEG during meditation and sleep in a personal environment?, 2) What EEG representations are better suited for prediction? and 3) How variable are the brain age predictions over long periods of time?

## 2 Methods

In this section, we describe how we trained machine learning models to predict age from low-cost mobile EEG and how the resulting brain age metric was evaluated.

### 2.1 Problem setting

We train a machine learning model *f*_**Θ**_ with parameters **Θ** to predict the age *y*^(*i*)^ of a subject *i* given their EEG ***X***^(*i*)^ *∈* ℝ^*C×T*^, with *C* the number of EEG channels and *T* the number of time points. To do so, *f*_**Θ**_ is trained to minimize the error between the true target age *y*^(*i*)^ and the predicted age *ŷ*^(*i*)^ over a training set of *N* subjects, *i*.*e*., *i ∈* [1, *N*]. Here, we use a Ridge regression model (Hoerl and Kennard, 1970) as *f*_**Θ**_. Following previous research on EEG-based modelling for brain age prediction, *f*_**Θ**_ is fed the output of a feature extractor ***ϕ*(*X*)** *∈* ℝ^*F*^, with *F* the number of features, which extracts multiple frequency-specific covariance matrices vectorized with Riemannian geometry (Sabbagh et al., 2019, 2020). Another viable option could have involved representations obtained through deep learning (Banville et al., 2021, 2022; Engemann et al., 2022; Roy et al., 2019).

### 2.2 Datasets

We used real-world mobile EEG recordings collected by anonymized users of the Muse S headband (RRID:SCR 014418; InteraXon Inc., Toronto, Canada). This data was collected in accordance with the privacy policy (July 2020) users agree to when using the Muse headband^1^ and which ensured their informed consent concerning the use of EEG data for scientific research purposes. The Muse S is a wireless, dry EEG device (TP9, Fp1, Fp2, TP10, referenced to Fpz) with a sampling rate of 256 Hz. Two types of recordings were used in this study: 1) meditation recordings with neurofeedback and 2) overnight sleep recordings. In both cases, recordings were carried out by users through the *Muse* application on iOS or Android mobile devices.^2^ In meditation recordings, users performed an eyes-closed focused attention task with real-time auditory feedback. This feedback was driven by the users’ EEG using a proprietary machine learning-based algorithm aimed at helping them focus on their breath. In overnight sleep recordings, users had the option to listen to various audio content and/or auditory feedback while falling asleep.

We collected four datasets by selecting recordings from InteraXon Inc.’s anonymized database of Muse users (see Table 1). First, the *Muse Meditation Dataset* (MMD), a subset of 971 meditation recordings from unique individuals, was obtained by sampling recordings of at least five minutes, with excellent signal quality, such that the age distribution was approximately uniform between 18 and 81 years of age (see Supplementary materials A for the detailed procedure).

**Table 1:**
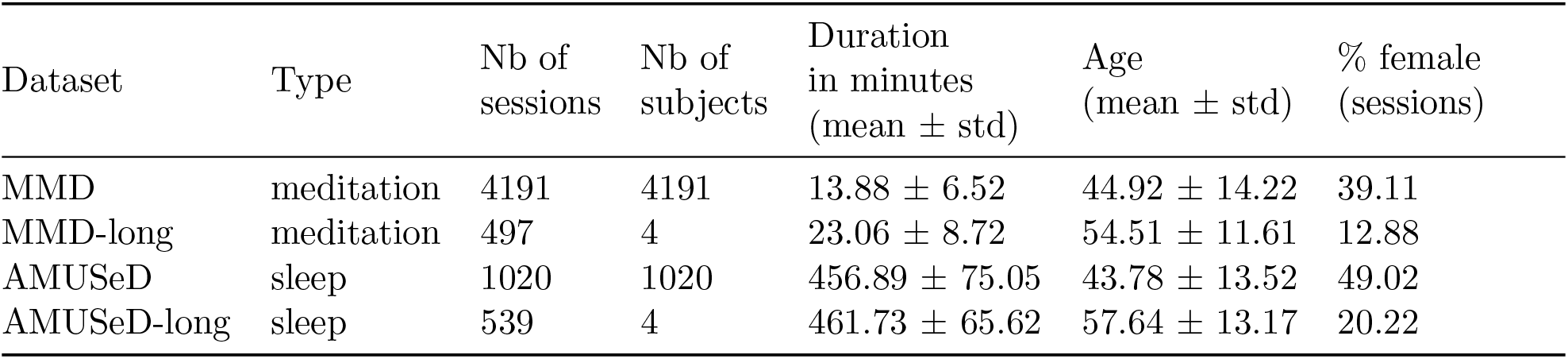
Description of the datasets used in this study.

Next, the *At-home Muse Unlabelled Sleep Dataset* (AMUSeD), a subset of 1020 overnight sleep recordings from unique individuals, was obtained in a similar fashion to MMD, *i*.*e*., we sampled sleep recordings of excellent signal quality, between five and 11 hours long, and with as uniform of an age distribution as possible were (see Supplementary materials B for more details). To study the long-term dynamics of EEG-based brain age, we additionally curated datasets containing sessions from a few users with a high number of recordings over multiple consecutive weeks and months. Following the same signal quality criteria as for MMD and AMUSeD, we sampled the *Longitudinal Muse Meditation Dataset* (MMD-long) and *Longitudinal At-home Muse Unlabelled Sleep Dataset* (AMUSeD-long), each containing recordings from four subjects (distinct subjects in both datasets) of different age groups. This yielded a total of 497 recordings for MMD-long and 539 recordings for AMUSeD-long^3^.

### 2.3 Preprocessing

Minimal preprocessing steps were applied to EEG data before feeding it to the prediction models. In the case of meditation recordings, the first minute of the meditation exercise (in which signal quality might still be settling) was cropped, and as much as eight of the following minutes of data were kept, depending on the length of the recording. Sleep recordings were kept in their entirety. Next, missing values (which can occur if the wireless connection is weak and Bluetooth packets are lost) were replaced through linear interpolation using surrounding valid samples.

A zero-phase FIR band-pass filter between 0.1 and 49 Hz was applied to both data types (Engemann et al., 2022). We additionally applied FIR notch filters to attenuate some hardware-specific spectral peaks that appeared in some of the recordings (at 16, 21.3, 32 and 42.7 Hz) as well as power line noise frequencies (50 and 60 Hz). Though conservative, this approach to noise reduction is simple to implement and led to improved brain age prediction performance.

Non-overlapping windows were then extracted: 10-s and 30-s windows were used for meditation and sleep recordings, respectively. In both cases, windows for which peak-to-peak amplitude exceeded a value of 250 *µ*V were rejected. Finally, EEG time series were downsampled to 128 Hz.

### 2.4 Age prediction pipelines

We relied on a commonly used method for EEG-based regression tasks, *i*.*e*., Ridge regression on filterbank covariance matrices with Riemannian geometry (Congedo et al., 2017; Engemann et al., 2022; Sabbagh et al., 2020) (see Supplementary materials D for more details). Each recording was represented by covariance matrices containing information from nine frequency bands estimated on the extracted non-overlapping windows. The covariance matrices were then vectorized and fed to the Ridge regression model.

To train models on sleep recordings, this approach was slightly modified such that one set of covariance matrices was extracted per sleep stage. By representing each sleep stage by its own set of features, we ensured that stage-specific spectral and spatial characteristics were made available to the machine learning models. We used an automatic sleep staging model (Abou Jaoude et al., 2020) trained on a labelled subset of Muse S overnight recordings to obtain sleep stage predictions for each 30-s window of AMUSeD and AMUSeD-long recordings (see Supplementary materials C for more details). Filterbank covariance matrices were then extracted per sleep stage following the same procedure as described above.

Models were trained on MMD or AMUSeD using Monte Carlo (shuffle split) cross-validation with 100 splits and 10% testing data. The training folds were further split to perform a hyperparameter search using generalized leave-one-out cross-validation for selecting the regularization strength. The regularization parameter was chosen among 100 values spaced log-uniformly between 10^*−*5^ and 10^10^. The examples from each subject were always used in only one of the training, validation or testing sets.

Predictions on the test folds were then used to evaluate the performance of the different models, as measured with the coefficient of determination *R*^2^ (Hughes and Grawoig, 1971), *i*.*e*., the percentage of chronological age variance explained by brain age.

Finally, models were retrained using the entire MMD or AMUSeD as training set (with the same validation set splitting strategies) and evaluated on the longitudinal recordings of MMD-long or AMUSeD-long.

## 3. Results

### 3.1 Predicting age from at-home mobile EEG recordings

Can low-cost, at-home mobile EEG be used to predict someone’s age, even if EEG is recorded with very few channels in environments that are not experimentally controlled? To explore this question, we trained machine learning models to predict age from such at-home EEG data on thousands of meditation or overnight sleep recordings. The performance obtained by these models is shown in Fig. 1. Scatter plots showing the relationship between chronological age and predicted brain age are shown in Fig. S1.

**Figure 1:**
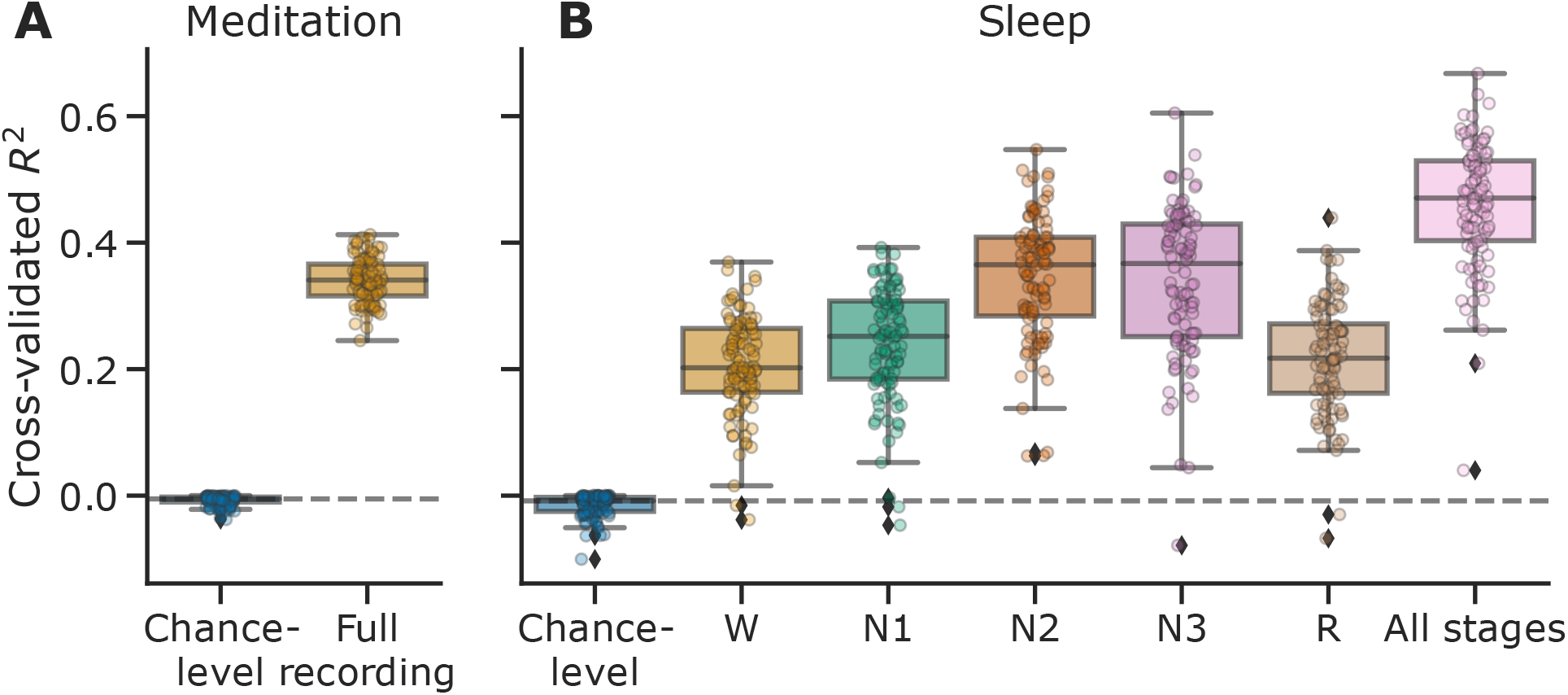
Cross-validated brain age prediction performance (*R*^2^) of models trained on at-home meditation or overnight sleep mobile EEG recordings (Monte Carlo, 100 iterations, 10% testing data). Points represent the scores of individual models trained on different cross-validation splits, while the boxplots display the overall score distributions. **A**. Models trained and evaluated on 4191 4- to 8-minute recordings of an eyes closed meditation exercise performed better than chance. **B**. Individual models trained on a dataset of 1020 overnight sleep recordings, using a single sleep stage (W, N1, N2, N3 or R) or all stages at once (“All stages”) also performed better than chance. See Fig. S2 for the performance of the same models reported as mean absolute error (MAE).

Models trained on meditation data (Fig. 1, left) achieved a median *R*^2^ of 0.34, markedly above chance (median of −0.01, as estimated by evaluating a dummy regressor which predicts the median age in the training set). Similarly, on sleep data (Fig. 1, right), all models achieved above-chance performance, with models trained on all sleep stages obtaining the highest performance (median *R*^2^ of 0.47 vs. −0.01 chance-level). The performance of models trained on either W or R stages alone was the lowest (0.20 and 0.22), while N2- and N3-based models both achieved higher performance (both 0.37). Performance was highest when combining information across all sleep stages, suggesting that additive information was conveyed by different sleep stages. Does this finding imply that sleep is even better suited for age prediction than meditation? While the results do show higher *R*^2^ for sleep-based models, the values cannot be directly compared as the age distribution, recording duration and sample sizes of the meditation and sleep datasets are not the same (as shown by different chance-level mean absolute error distributions, see Fig. S2). In sum, these findings show that machine learning models initially developed for the laboratory setting can accurately predict an individual’s age based on their EEG collected with an at-home mobile device from either sleep or meditation contexts.

### 3.2 Inspection of trained brain age models

What drives the predictions of these models, *i*.*e*., what kind of information is critical for predicting age? Knowing this would not only validate the functioning of our models, but also provide interesting information to identify markers of aging in the human brain. To answer this question, we trained additional brain age models on the meditation and sleep datasets, but this time with inputs that capture different types of information. Results are presented in Fig. 2.

**Figure 2:**
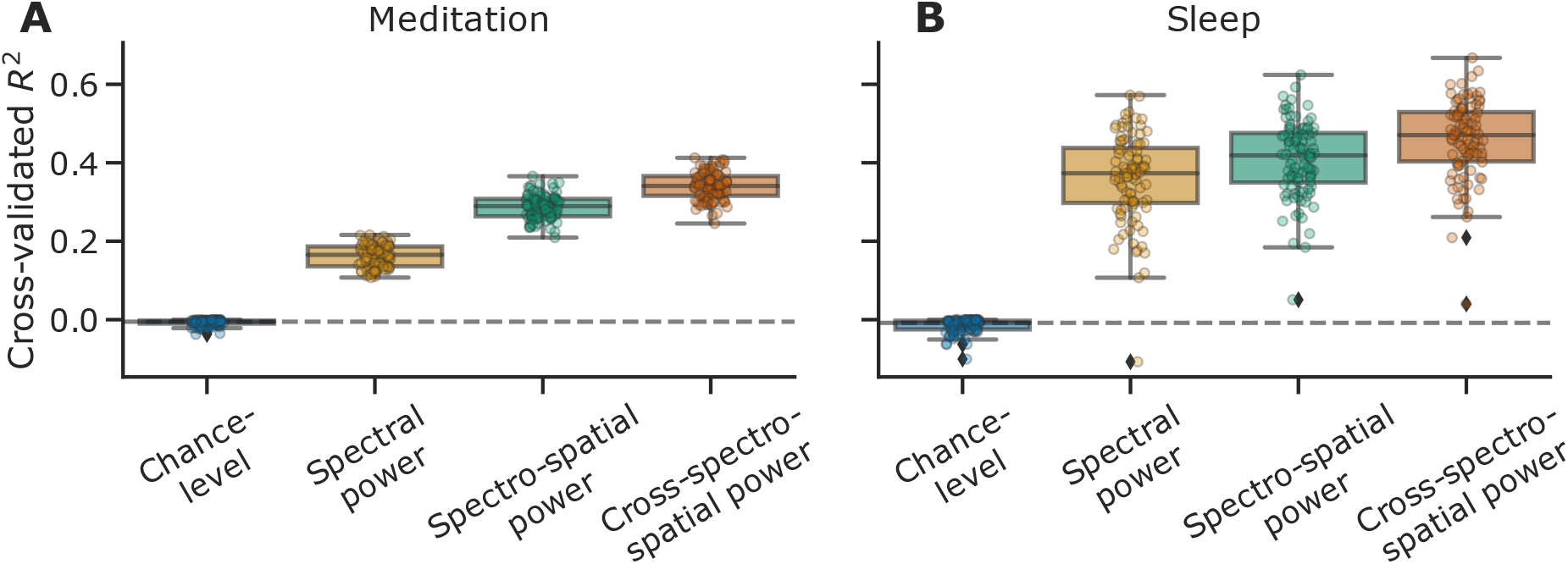
Cross-validated brain age prediction performance (*R*^2^) of models trained on different EEG input representations (Monte Carlo, 100 iterations, 10% testing data) for (**A**) meditation and (**B**) sleep recordings. For sleep results, models based on all sleep stages (“All stages” in Fig. 1) were used. For both meditation and sleep data, the inclusion of spectral, spatial and cross-spectral information yielded the best performance despite the limited number of only four EEG channels.

For both meditation- and sleep-based models, providing spectral information only (*i*.*e*., frequency band log-powers for each EEG channel) already yielded performance visibly above chance. Adding further spatial information, *i*.*e*., by allowing the model to learn from the correlation between pairs of channels, yielded a clear improvement for both data types, with 100/100 and 85/100 of the trained “spectro-spatial” models reaching better performance across cross-validation iterations than their equivalent “spectral”-only models for meditation and sleep data, respectively. This suggests that spatial information might be slightly more useful for decoding age from meditation rather than from sleep data. Finally, additionally allowing the models to learn from cross-frequency correlations helped improve performance for both meditation and sleep models, *i*.*e*., cross-spectro-spatial models were better than spectro-spatial models on 99/100 and 96/100 cross-validation iterations, respectively.

Given that strictly spectral-based models already yielded above-chance age prediction performance, we can further inspect them to see which frequency bands provided the most useful information. As shown by a permutation importance analysis^4^ (Breiman, 2001), different bands were useful for the meditation- and sleep-based models (Fig. 3). On meditation data, low and mid *α* sub-bands (8-10 and 10-12 Hz) were the most important, *i*.*e*., making this information unusable by the models resulted in the largest performance decrease. On sleep data, given that the spectral information was extracted per sleep stage, it is possible to consider the importance of the different bands in each sleep stage (Fig. 3, right). Strikingly, the most useful feature was *α*_*high*_ (12-15 Hz) in N2, whose importance was about twice as high as the second most useful feature, *δ;* power (1-4 Hz) in N3. Moreover, mid and high *α* (10-12 and 12-15 Hz) in N3 also stood out as critical features. The frequency ranges and sleep stages identified above relate to well-known characteristics of awake and sleep EEG (*e*.*g*., alpha peak, sleep spindles and slow waves), suggesting that the models picked up biological information.

**Figure 3:**
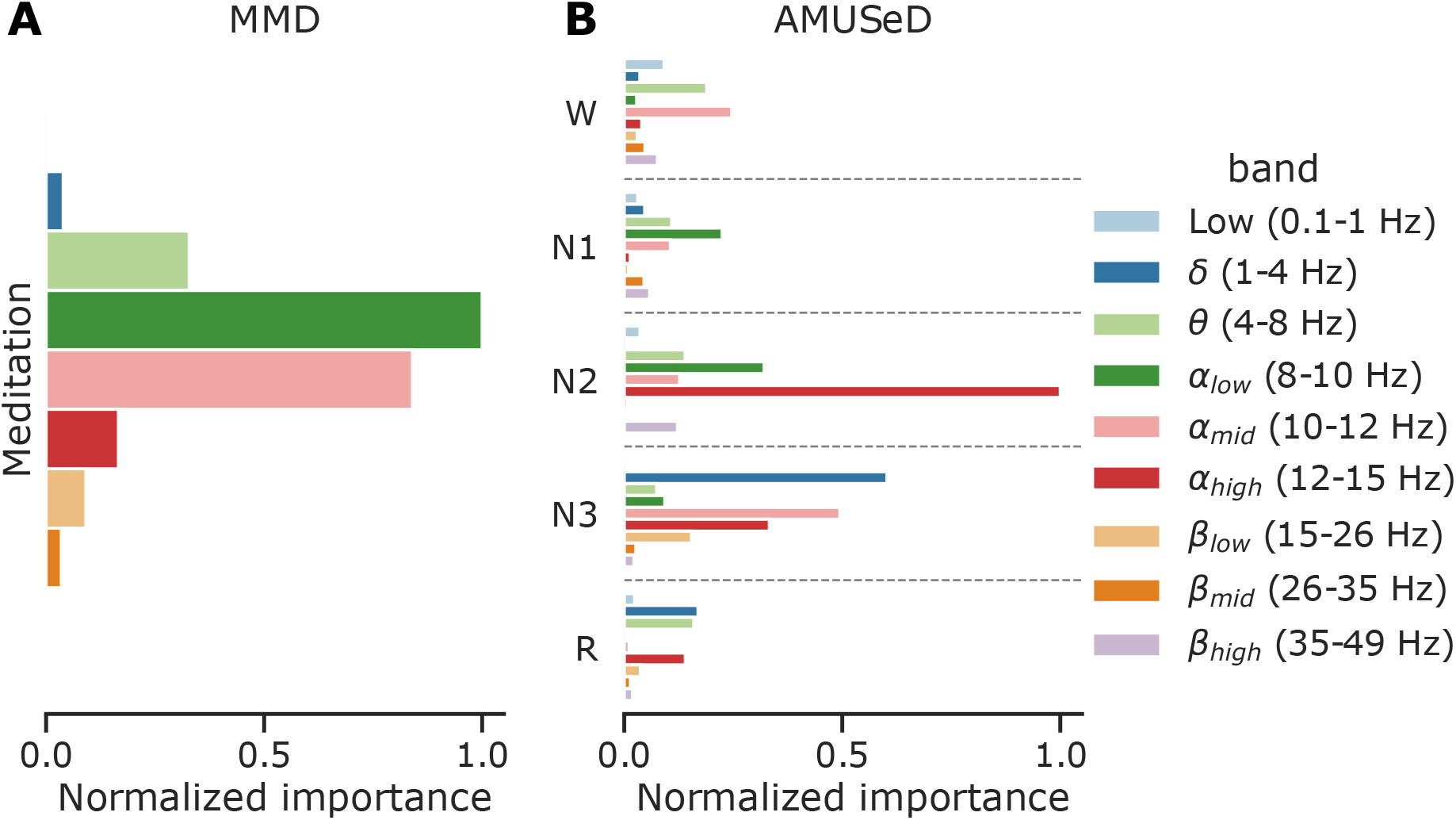
Permutation analysis using the spectral power models trained on (**A**) meditation and (**B**) sleep data. The x-axis indicates the average performance variation (∆*R*^2^, normalized between 0 and 1 for comparability) obtained when the values in a specific frequency band or a combination of a frequency band and sleep stage are randomly permuted over 100 repetitions at testing time. On meditation data, models were most sensitive to *α*_low_ and *α*_mid_ bands, while on sleep data models were most sensitive to *α*_high_ in N2 and *δ;* and *α*_mid_ in N3.

Overall, these results suggest that spectral, spatial and cross-spectral dimensions all provide complementary information that can help predict an individual’s age from their mobile EEG. By evaluating the importance of the different frequency bands, we further demonstrated that our models relied on specific EEG patterns to predict age, providing initial validation of our approach.

### 3.3 Variability of brain age over multiple weeks and months

To assess the practicality of the brain age metric obtained from home-based EEG with few electrodes, it is helpful to consider its variability over medium- to long-term periods. Indeed, stable values over weeks and months would indicate actual “trait”-level information, *e*.*g*., related to aging and, potentially, pathological aging, is being captured. On the other hand, significant variability at smaller scales would suggest the metric is also influenced by momentary factors and therefore captures “brain states” as well. To answer these questions, we computed the recording-by-recording brain age of a few subjects for whom a large number of recordings was available and inspected their predicted brain age ∆ values. The results of this analysis are presented in Fig. 4 (see Fig. S4 for additional results).

**Figure 4:**
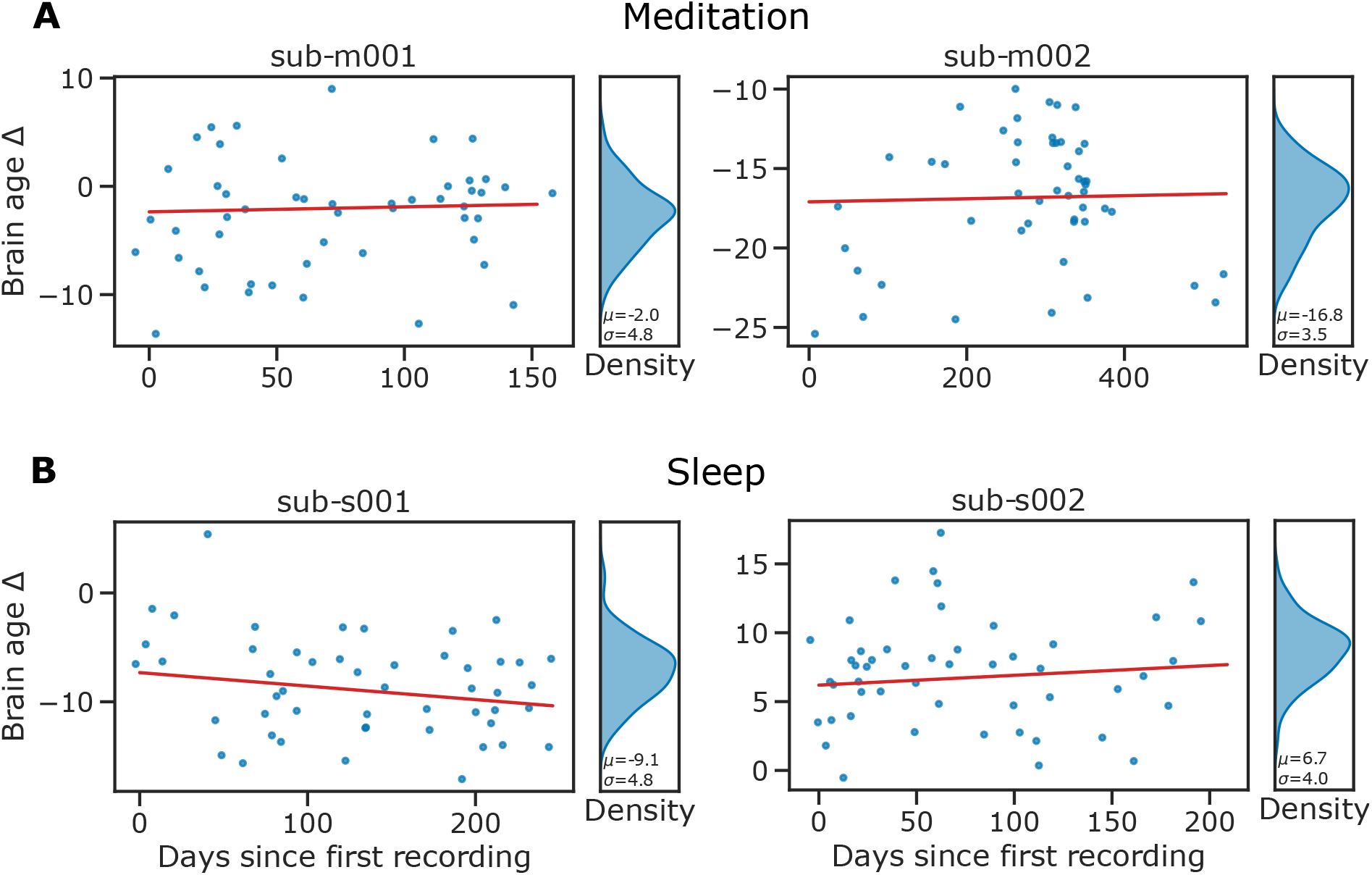
Longitudinal brain age ∆ predictions for four subjects with (**A**) multiple consecutive meditation or (**B**) sleep recordings. Four subjects with more than 50 recordings of good signal quality were selected. Their brain age was predicted using a model trained on MMD or AMUSeD. Blue dots represent the brain age ∆ predicted for single recordings. To ensure the anonymity of the subjects, we only presented 50 randomly sampled recordings for each subject and added random jitter *δ; ~ N* (0, 20 days) to the recording dates. Nevertheless, all available sessions were used to fit a linear model (red line) showing the trend for each subject. Density plots summarize the distribution of predicted age (blue marginal plots). Despite significant variability visible in all subjects, the slopes obtained from linear models across sessions (red lines) were close to zero, pointing at a stable average brain age across time. This suggests the proposed brain age metric captures both “trait”- and “state”-like information.

While there is substantial longitudinal variability across recordings from a same subject, the average predicted values remain fairly stable across longer-term periods, for both meditation- and sleep-based models. Despite across-recording standard deviations of 3.5 to 6.0 for meditation, and 3.3 to 5.5 for sleep, the average predictions over longer periods (*e*.*g*., months) do not vary substantially as can be seen from the relatively flat slopes obtained from linear models (min = *−*0.017, max = 0.015). This is seen both in subjects whose chronological and brain age match closely (*e*.*g*., sub-m001) and in subjects for whom there is a larger difference between the two measures (*e*.*g*., sub-m002).

Taken together, these results suggest that our brain age metric derived from mobile EEG captures “trait”-like information, but also shorter-term “state”-like information which may reflect a subject’s state at the time of the recording.

## 4. Discussion

In this paper, we showed that brain age measurement, thus far restricted to clinical and laboratory settings, can be extended to at-home settings using low-cost mobile EEG. We presented experiments on over 5200 recordings from unique individuals, combining real-world data collected during meditation exercises and overnight sleep. Results highlighted the usefulness of cross-spatio-spectral information to predict age, and revealed how certain EEG characteristics, such as *α*_*low*_ power for meditation, N2 *α*_*high*_ power in the spindle range and N3 *δ;* power are important for accurately predicting age. Inspired by how mobile EEG technology could facilitate longitudinal monitoring of brain biomarkers, we investigated the stability of brain age over repeated sessions spanning a few months to over a year for eight individuals. This exploration revealed short-term recording-to-recording variability but also stability across longer-term periods that have not been reported in the previous literature focusing on very few or single snapshots of individual EEG (Banville et al., 2021; Engemann et al., 2022; Sabbagh et al., 2020; Sun and Saenko, 2016). Our results provide a foundation for the development of brain age assessment tools that can be applied out-of-the-lab with real-world EEG.

### 4.1 Biomarker measurement in real-world conditions with mobile EEG

The results presented in this work were obtained on recordings carried out at-home by users themselves, without expert supervision or a complicated experimental setup, relying instead on off-the-shelf, low-cost mobile EEG hardware. Because it is both an easier and cheaper way to record data, this approach could enable collecting larger and more diverse samples. Concretely, we trained models on datasets containing more than 5200 of at-home EEG recordings (see Table 1). This is an order of magnitude above much of the published work on EEG-based brain age and in line with the previous studies with the highest sample sizes (Brink-Kjaer et al., 2022; Engemann et al., 2022; Sun et al., 2019). Critically, these studies focused on clinical or research datasets which are much more expensive and time-consuming to collect. Despite working on noisier real-world EEG data, brain age prediction performance was in a similar range as these published results, *e*.*g*., 0.47 vs. 0.33-0.61 *R*^2^ in Engemann et al. (2022) on their largest reported dataset, and 7.6 (Fig. S2, “All stages”) vs. 7.8 years mean absolute error in Sun et al. (2019). Likely, this similarity in performance reflects the fact that sleep EEG is a rich source of physiological information and that a few channels are sufficient to capture some of its age-related features well. Moreover, our results support the use of a prediction pipeline based on filterbank covariance matrices and Riemannian geometry, as previously proposed for brain age modelling (Engemann et al., 2022; Sabbagh et al., 2020, 2022).

The ease-of-use and low friction provided by mobile EEG devices also make it easier to collect longitudinal data over extended periods of time. This has allowed us to evaluate brain age longitudinally on an unprecedented number of repeat sessions, *i*.*e*., 1036 recordings from four subjects spanning periods of up to 529 days, while previous work was limited to a few days, *e*.*g*., four recordings collected over consecutive days or a few days apart (Hogan et al., 2021). This highlights the potential of mobile EEG to bring biomarker monitoring out-of-the-lab and into the real world.

### 4.2 What does EEG-based brain age capture?

Brain age models trained on structural MRI data rely on anatomical features such as cortical thickness and brain volume to provide accurate predictions of one’s chronological age (Cole et al., 2019). As EEG captures brain activity, rather than anatomy, what specific information does it contain that enables age prediction? Given the important differences between awake and sleep EEG, the answer differs for the types of data presented in this study.

As shown in Section 3.1, sleep EEG provided higher age prediction performance than awake EEG. This was not entirely surprising, as there is a well-known connection between sleep microstructure and age (Muehlroth and Werkle-Bergner, 2020; Purcell et al., 2017), meaning the predictive models have access to rich age-related information (through the cross-spectro-spatial representation described in Section 2.4). This is in line with Section 3.2 which revealed that the most useful features for predicting age are in fact descriptors of key sleep microstructure components. For instance, *α*_*high*_ (12-15 Hz) power in N2, the most important feature according to the analysis, fits with the *σ* band (11-15 Hz) sleep spindles typically fall in (Purcell et al., 2017). The second most useful feature, *δ;* (1-4 Hz) power in N3, captures the well-documented age-related decrease in slow wave (*<*4 Hz) power (Muehlroth and Werkle-Bergner, 2020). Nonetheless, combining information from all sleep stages yielded the best age prediction performance (see Fig. 1), suggesting there is additional useful information that might not be strictly related to well-known microevents. Similarly, including data from the entire night, rather than from the beginning or the end, leads to better performing models, as seen in the supplementary analysis of Fig. S3. These results confirm overnight sleep EEG is a rich source of information for modeling brain age.

Despite lower performance, models based on awake meditation EEG also yielded reasonable brain age predictions. Previous work on very similar cross-sectional data but from a descriptive, rather than predictive, point-of-view has shown that the characteristics of the *α* band vary significantly throughout adulthood (Hashemi et al., 2016). For instance, the *α* peak frequency decreases with age (a widely reported finding, see *e*.*g*., Chiang et al. (2011); Klimesch (1999); *Knyazeva et al. (2018); Merkin et al. (2023)). Additionally, a small but significant increase in α* and *β* power with age has also been reported in conjunction with a decrease in *δ;* and *θ* power (Hashemi et al., 2016). Indeed, our filterbank models heavily relied on the information contained in the low- and mid-*α* bands, *i*.*e*., between 8 and 12 Hz (see Fig. 3). As shown in the supplementary analysis of Fig. S5, visualization of the spectrum around the *α* band similarly confirmed that the peak *α* frequency decreased across age groups.

Overall, brain age models based on mobile EEG can capture known age-related electrophysiological phenomena. This provides early support for extending brain age measures beyond anatomical and lab-based assessments.

### 4.3 What causes brain age variability?

Longitudinal monitoring on eight subjects showed that while the average brain age ∆ predictions were mostly stable across longer periods of time (*e*.*g*., months), there was significant variability from one day to the next. Of note, another study which looked at EEG-based brain age across days, though on a shorter temporal scale, found a night-to-night standard deviation in brain age ∆ of 7.5 years (Hogan et al., 2021). This standard deviation could be further decreased to 3 years when averaging four nights. In comparison, our experiments of Section 3.3 on sleep data showed a standard deviation of 3.3 to 5.5 years for four subjects with at least 50 recordings each. While additional experiments on more subjects are required to validate these results, this suggests that similar variability is observed even with longer sleep EEG recordings. While a supplementary analysis (Supplementary materials G) suggests that time of day, at least cross-sectionally, is negatively correlated with brain age ∆ (Pearson’s *ρ* = *−*0.079, *p <* 10^*−*5^), most of the variability remained unexplained. What does this residual variability reflect?

On sleep data, intrinsic factors, *e*.*g*., fluctuations in sleep quality, likely account for some of this variability. For instance, increased slow wave activity (Plante et al., 2016) and decreased sleep spindle density but increased amplitude (Knoblauch et al., 2003) are established effects of sleep deprivation. As our models rely on *δ;* power in N3 to predict age and *σ* power in N2, the fluctuations might in fact partially reflect sleep quality or even past sleep dept. Similarly, for awake meditation data, different factors are known to influence the *α* peak, whose characteristics were most important for predicting age in our sample. For instance, research on “mental fatigue” has revealed that the spectral properties of EEG change as fatigue increases with, for instance, power in *θ, α* and *β* bands increasing (Arnau et al., 2017). Multiple studies on meditation and EEG also reported decreased *α* peak frequency and increased *α* power during meditation exercises (Cahn and Polich, 2006; Lomas et al., 2015). The observed variability might therefore reflect the quality of the meditation, the specific type of meditation exercise carried out during the recording or overall well-being related to recovery from activities during the day before the recording.

The observed patterns might appear inconsistent with the expected stability of an age-related measure, *i*.*e*., to be evolving slowly in time. However, it is important to recall that because existing brain age studies are typically limited to single or very few recordings per subject, similar fluctuations between recordings may simply be unknown. On the other hand, one could hypothesize that high-density EEG and a focus on EEG features beyond spectral power, *e*.*g*., measures of connectivity, may improve re-test reliability (Rolle et al., 2022). Unless head-to-head comparisons between mobile devices and laboratory-grade EEG are performed, this remains speculative. But in case future studies will uncover strong intrinsic variability between recordings across device types, mobile EEG assessments would have a clear advantage. Performing multiple repeat measures to estimate trait-like signatures will be far more tractable with home-based EEG.

### 4.4 Limitations

The datasets used in our experiments were collected in uncontrolled, at-home conditions, *i*.*e*., no expert monitoring was available to ensure compliance with the recording instructions. This contrasts with existing work on brain age prediction, where datasets are usually collected in clinical or research settings. Similarly, only minimal metadata was available in the datasets. As a result, the datasets did not contain the necessary information to separate healthy individuals (used for training) from pathological individuals (used for testing), as is commonly done in brain age studies. Since our models were likely exposed to some pathological data during training, their sensitivity to pathology-related EEG activity might be lower than in previous studies realized in clinical settings. Likewise, we did not have access to clinical cognitive assessments that would have allowed exploration and validation of brain age semantics, e.g., relating positive brain age ∆ to cognitive slowing (Denissen et al., 2022; Engemann et al., 2020). Finally, the datasets were not collected under controlled conditions typical of laboratory- or clinical-based studies, leaving open the possibility that individuals did not enter their actual age when creating their user profile. However, thanks to the type of sample size enabled by mobile EEG devices, much larger than traditional studies, our models will be less sensitive to the occasional incorrect label. For instance, a study using similar datasets produced results that agreed with earlier lab-based studies (Hashemi et al., 2016), supporting the validity of our dataset for brain age analysis.

## 5 Conclusion

Despite the challenges that come with out-of-the-lab collection of EEG, individual physiological characteristics can be successfully modeled during sleep and meditation using state-of-the-art machine learning. Brain age prediction, in particular, is a promising framework to leverage large-scale real-world EEG data. Combined with these tools and thanks to its cost-effectiveness and ease-of-use, at-home low-cost mobile EEG offers a promising opportunity to democratize, accelerate and scale up biomarker development.

## Acknowledgements

This work was supported by Mitacs (project number IT14765) and InteraXon Inc. (graduate funding support) for HB and by the ANR BrAIN AI chair (ANR-20-CHIA-0016) and ANR AI-Cog (ANR-20-IADJ-0002) grants for AG and DAE.

## Author contributions

Conceptualization: DAE, AG. Data curation: HB. Software: HB, DAE, AG, MAJ. Supervision: AG, DAE, CA. Investigation: HB. Visualization: HB, DAE. Methodology: AG, DAE. Writing: HB, DAE, AG, SUNW, SCH.

## Footnotes

1 https://choosemuse.com/legal/privacy/

2 https://choosemuse.com/muse-app/

3 We do not disclose these subjects’ ages or gender in order to minimize identifiable information.

4 The permutation analysis measures how the model’s performance, measured as *R*^2^, changes when one of its features has been randomly shuffled across the examples of the test set. Here, we normalized the ∆*R*^2^ values between 0 and 1 so that feature importance was straightforward to compare between the two datasets.

