## Supplementary materials for "Do try this at home: Age prediction from sleep and meditation with large-scale low-cost mobile EEG"

#### A Sampling of MMD

Meditation recordings of five minutes and more were selected from Muse S meditation recordings collected between October 2019 and October 2021. Recordings were then filtered to only keep users whose age was between 18 and 81 years at the time of recording. A single recording was sampled per user, such that the age distribution across all sampled recordings was approximately uniform and the dataset was balanced for male and female users. From this set, only recordings with excellent signal quality, based on basic signal statistics defined as follows, were retained. First, recordings for which more than 5% of samples were missing for any of the EEG channels (caused by Bluetooth packet loss during transmission from the headband to the mobile device) were rejected. The variance of the signals bandpass-filtered between 2 and 26 Hz was also computed for non-overlapping 1-s windows. Recordings for which, for any of the four channels, 25% or more of the windows had a variance above a threshold of 100 $\mu V^2$ were rejected. This finally yielded a subset of 4191 recordings (mean duration: 13.88 $\pm$ 6.52 minutes) with excellent signal quality from 4191 unique individuals. Mean age across recordings was 44.92 $\pm$ 14.22 years old (min: 18, max: 81) and 39.11% of recordings were of female users.

#### B Sampling of AMUSeD

We sampled sleep recordings, following a similar procedure to the one described above for MMD. Recordings that lasted between 5 and 11 hours, from users between 18 and 81 years of age at the time of recording, and which were started between 5 PM and 5 AM, local time, were selected. A maximum of two recordings per user were selected, which were then screened for signal quality. Signal quality screening was performed on non-overlapping 30-s windows (*i.e.,* the standard window length in sleep recording analysis). Recordings for which more than 25% of the windows had a variance above 1,000 $\mu V^2$ were rejected. Finally, the recording with the highest number of good windows was kept for each user. This yielded a total of 1020 overnight sleep recordings from 1020 unique users, with a mean duration of 456.89 $\pm$ 75.05 minutes, a mean age across recordings of 43.78 $\pm$ 13.52 years (min: 18, max: 79) and 20.22% of recordings of female users.

#### C Automatic sleep staging

In order to train brain age models on sleep stage-specific features, we obtained sleep stage predictions using a classifier trained on labeled Muse S data with classes W (wake), N1, N2,

N3 (non-REM sleep) and R (rapid eye movement sleep). A convolutional and recurrent neural network architecture inspired by Abou Jaoude et al. (2020) was trained on Muse S sleep recordings annotated according to the AASM guidelines (Berry et al., 2012) by a sleep technician (see Banville et al. (2022) for a description of the dataset). In contrast to the original architecture, the model 1) used unidirectional, rather than bidirectional, LSTM layers, to allow its use in a real-time processing context and 2) had different input layer and maxpooling kernel sizes, in order to accommodate signals sampled at 128 Hz (vs. 200 Hz in Abou Jaoude et al. (2020)). The model was trained to predict which sleep stage (W, N1, N2, N3 or R) a 30-s EEG window corresponded to. Moreover, the data was preprocessed in a similar manner to what is described in Section 2.3: 1) linear interpolation of missing values, 2) downsampling to 128 Hz, 3) bandpass filtering between 1 and 40 Hz, and 4) channel-wise zero-meaning of each window. The trained model was deployed as-is on the sleep recordings of AMUSeD and AMUSeD-long to produce hypnograms, *i.e.,* sequences of overnight sleep stage predictions, with 30-s resolution.

### D Detailed description of the age prediction pipeline

The filterbank covariance pipeline first applied a filter bank to the input EEG, yielding narrow-band signals in the nine following bands: low frequencies (0.1-1 Hz), $\delta$ (1-4 Hz), $\theta$ (4-8 Hz), $\alpha_{\text{low}}$ (8-10 Hz), $\alpha_{\text{mid}}$ (10-12 Hz), $\alpha_{\text{high}}$ (12-15 Hz), $\beta_{\text{low}}$ (15-26 Hz), $\beta_{\text{mid}}$ (26-35 Hz) and $\beta_{\text{high}}$ (35-49 Hz). Covariance matrices were estimated in each frequency band from non-overlapping 10-s window using the OAS algorithm (Chen et al., 2010), yielding a set of nine covariance matrices per recording. In a first variation , the log-diagonal of each covariance matrix was extracted and concatenated into a single feature vector of dimension $C \times 9$, z-score normalized using the mean and standard deviation of the training set, then fed to a Ridge regression model (the "spectral power" model in Fig. 2). For the "spectro-spatial power" variation, covariance matrices were instead projected into their Riemannian tangent space, exploiting the Wasserstein distance to estimate the mean covariance used as the reference point (Bhatia et al., 2018; Sabbagh et al., 2019). The vectorized covariance matrices with dimensionality $C(C+1)/2$ were finally z-score normalized, concatenated, and fed to a linear regression model with an $L_2$ penalty, just as for the "diag" model. Finally, for the "cross-spectro-spatial power" variation, the filtered signals in the nine frequency bands were concatenated and a single $C \times 9$ by $C \times 9$ covariance matrix was computed. This allowed further capturing interactions between channels in different frequency bands. The same vectorization and regression approach as for the "spectro-spatial" model above was then used but with this covariance matrix as input.

The pipeline was modified in sleep experiments to capture information about each sleep stage independently. To this effect, one set of covariance matrices was extracted and vectorized per sleep stage (following the three approaches described above). Features from all five stages were then concatenated and fed to a single Ridge regression model.

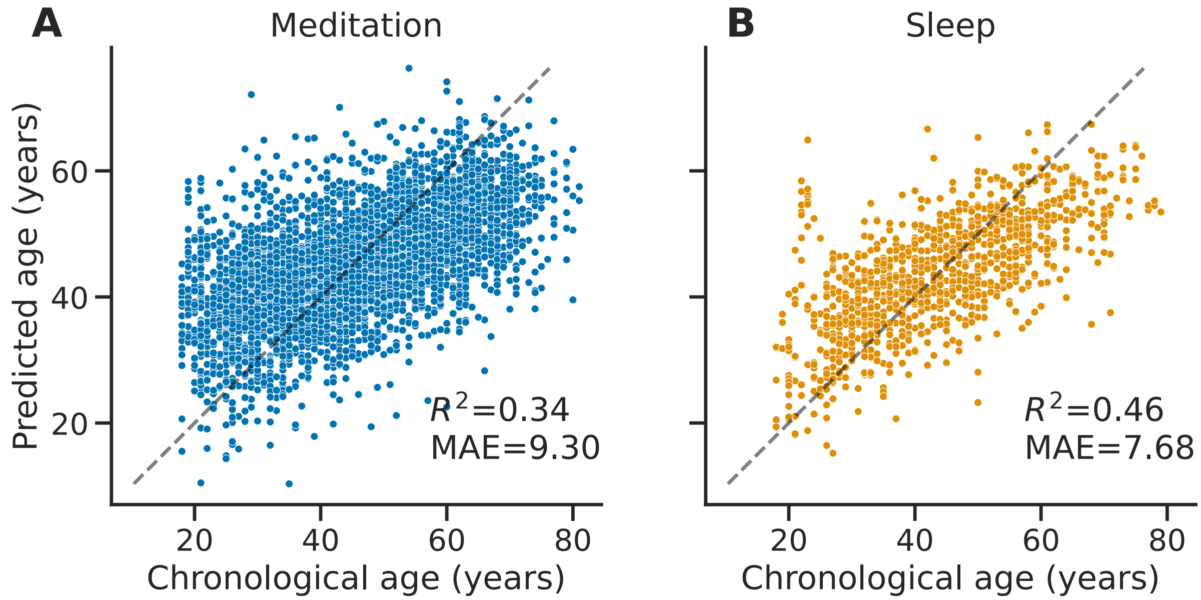


Figure S1: Scatter plots of chronological age versus predicted brain age for (**A**) meditation recordings and (**B**) sleep recordings. Cross-validated age predictions were obtained using cross-spectro-spatial power models trained over MMD or AMUSeD, respectively, with 10-fold cross-validation. $R^2$ and MAE are reported for the plotted predictions. Each point represents a single subject and the identity lines indicate the behavior of a perfect model.

A combination of the `MNE-Python` (Gramfort et al., 2014), `pyRiemann` (Barachant et al., 2013), `coffeine` (Sabbagh et al., 2020), `mne-features` (Schiratti et al., 2018), `scikit-learn` (Pedregosa et al., 2011), `mne-bids` (Appelhoff et al., 2019) and `mne-bids-pipeline` (Jas et al., 2018) packages were used to carry out our experiments.

#### E Additional brain age prediction results

Fig. S1 show the relationship between the brain age predictions obtained on MMD and AMUSeD and the true chronological age of each subject.

Fig. S2 presents the results of the brain age modeling experiments of Fig. 1 but using the mean absolute error instead of $R^2$. Fig. S3 presents the performance of brain age models trained on AMUSeD using different parts of the night. Fig. S4 shows longitudinal brain age estimates for four additional subjects.

#### F EEG spectral characteristics in MMD

A visual assessment of the $\alpha$ peak characteristics of the data in MMD reveals that both the $\alpha$ peak frequency, *i.e.,* the frequency of maximum power in the $\alpha$ band, as well as the $\alpha$ power,

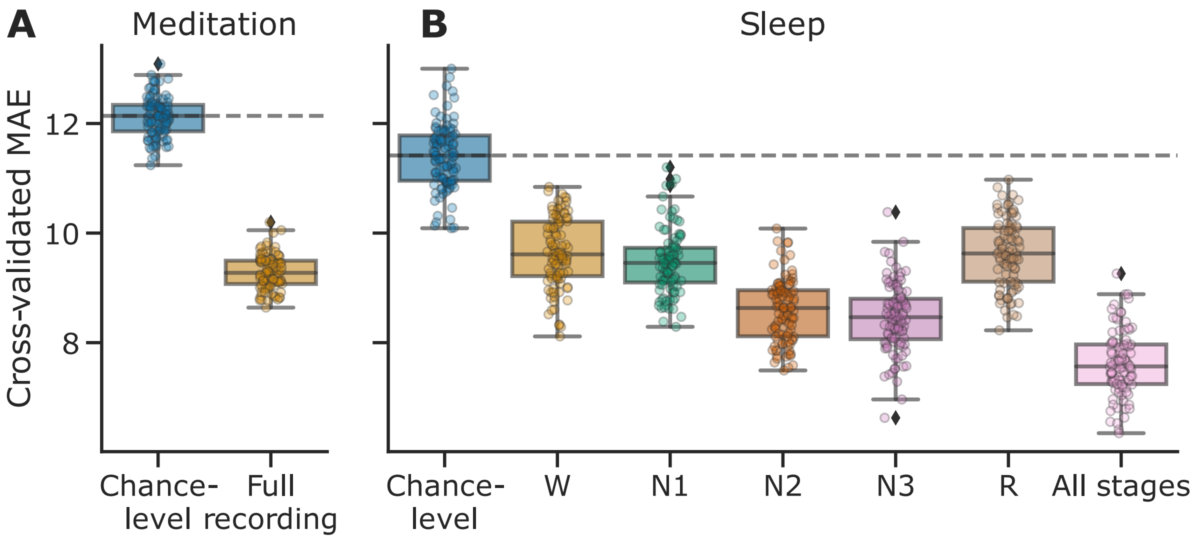


Figure S2: Cross-validated brain age prediction performance as measured with the mean absolute error (MAE). See Fig. 1 for a full description.

vary across age groups (see Fig. S5). Therefore, these features are likely useful for distinguishing age groups in our meditation datasets.

#### G Time of day effects on brain age $\Delta$

Fig. S6 shows brain age $\Delta$ measures for all recordings of the MMD dataset as a function of time of day. Brain age $\Delta$ and time of day are significantly correlated (Pearson's $\rho = -0.079$, $p < 10^{-5}$).

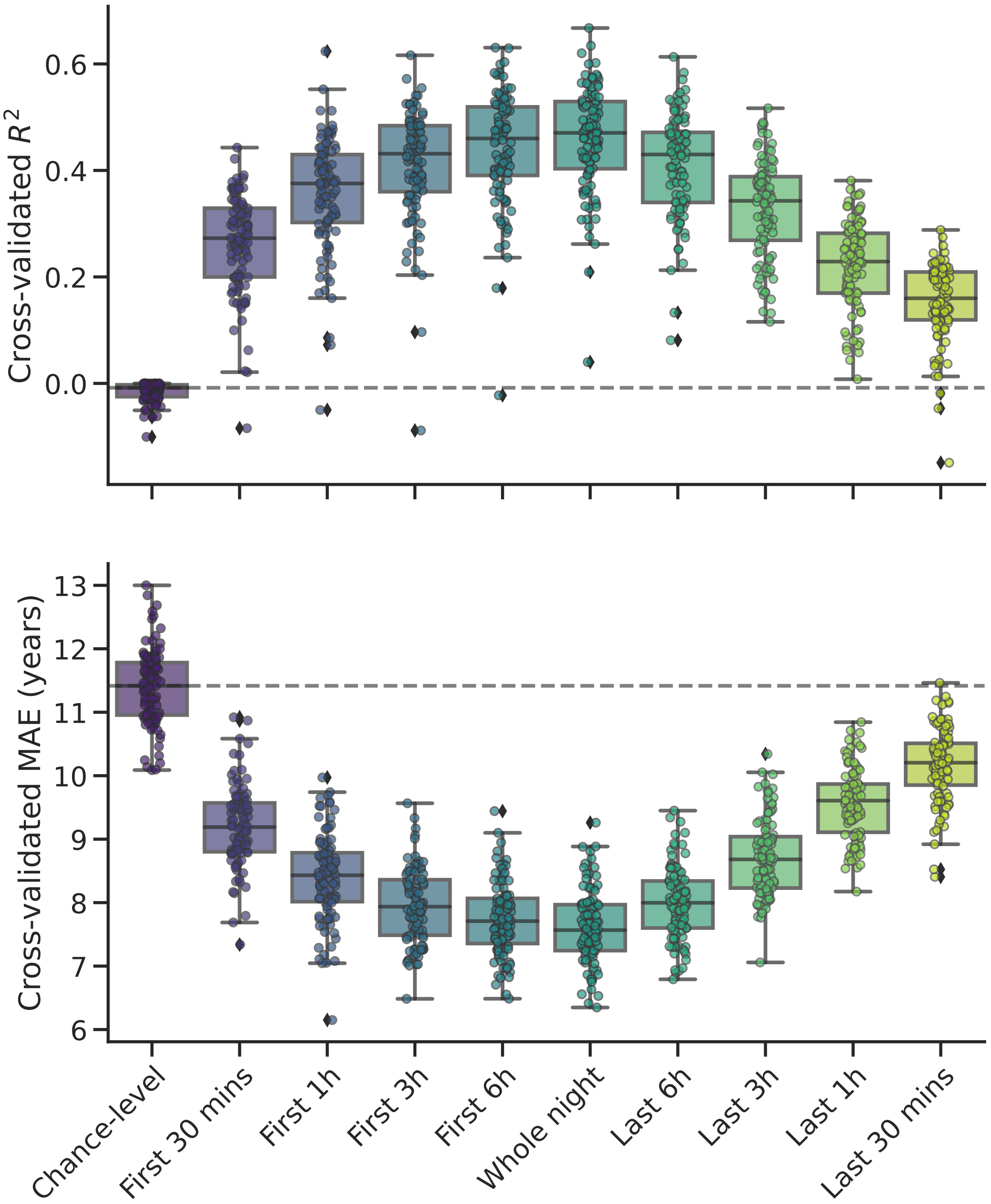


Figure S3: Cross-validated brain age prediction performance of models trained on different parts of overnight sleep recordings (Monte Carlo, 100 iterations, 10% testing data) measured with (**A**) $R^2$ and (**B**) MAE. See Fig. 1 for details. Separate models were trained on the first part of the night (first 30 mins, 1h, 3h or 6h) or the last part of the night (last 6h, 3h, 1h or 30 mins). While a longer duration generally helped performance, models based on the first part of the night performed better than those based on the last part, likely because it contains higher proportion of NREM sleep.

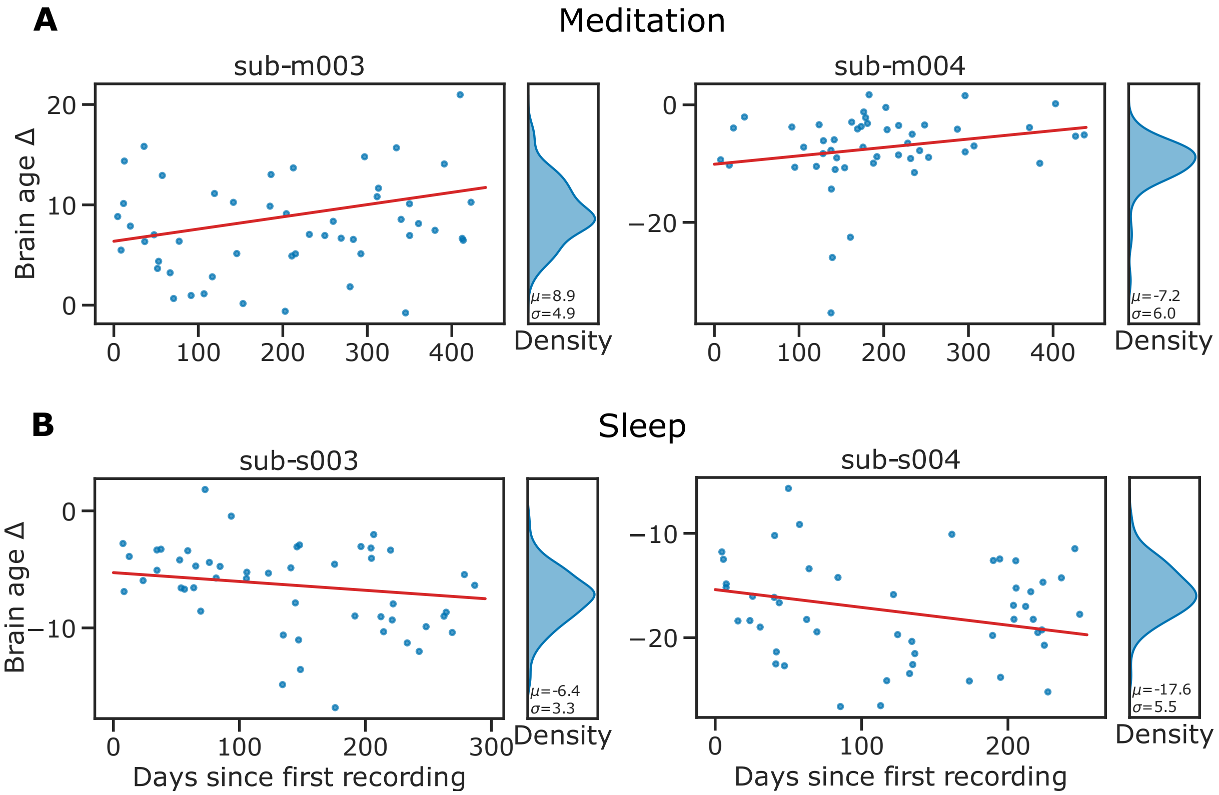


Figure S4: Longitudinal brain age $\Delta$ predictions for four additional subjects with multiple consecutive meditation (top) or sleep (bottom) recordings. See Fig. 4 for details.

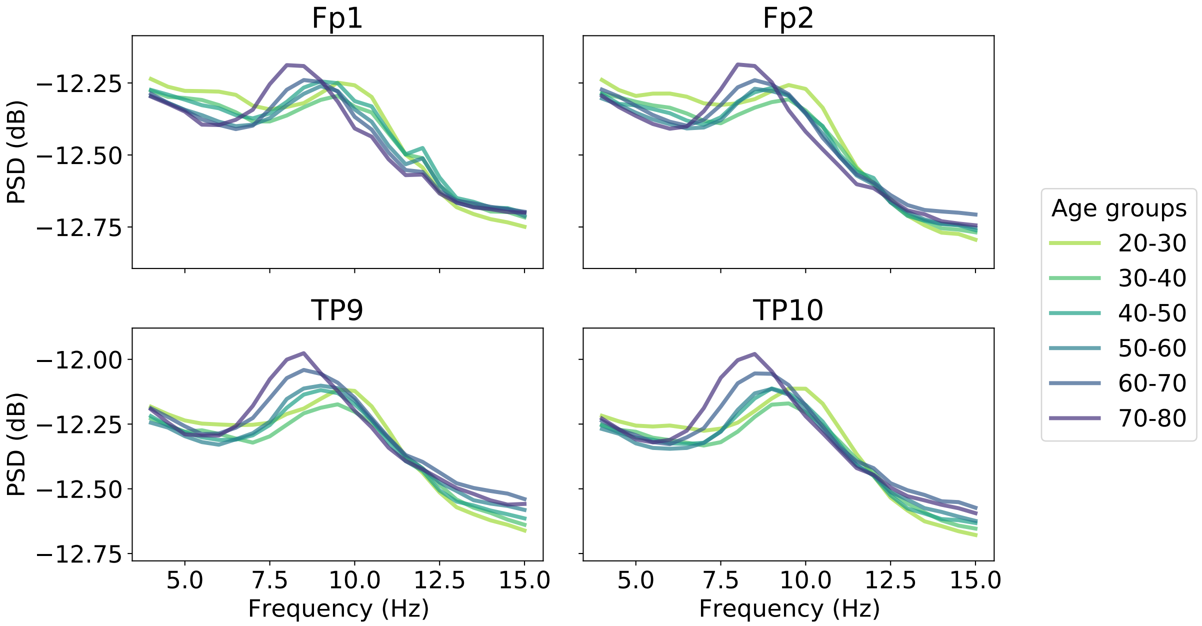


Figure S5: Visualization of the spectrum around the $\alpha$ band for recordings of MMD. Welch's periodogram with non-overlapping 4-s segments and median aggregation was applied to the 10-s windows preprocessed for use with filterbank models. The PSDs were then $\log_{10}$ −transformed and averaged inside each 10-year slice of the subjects. Older individuals tended to have a lower $\alpha$ peak frequency, as well as higher $\alpha$ peak power, in all four electrodes.

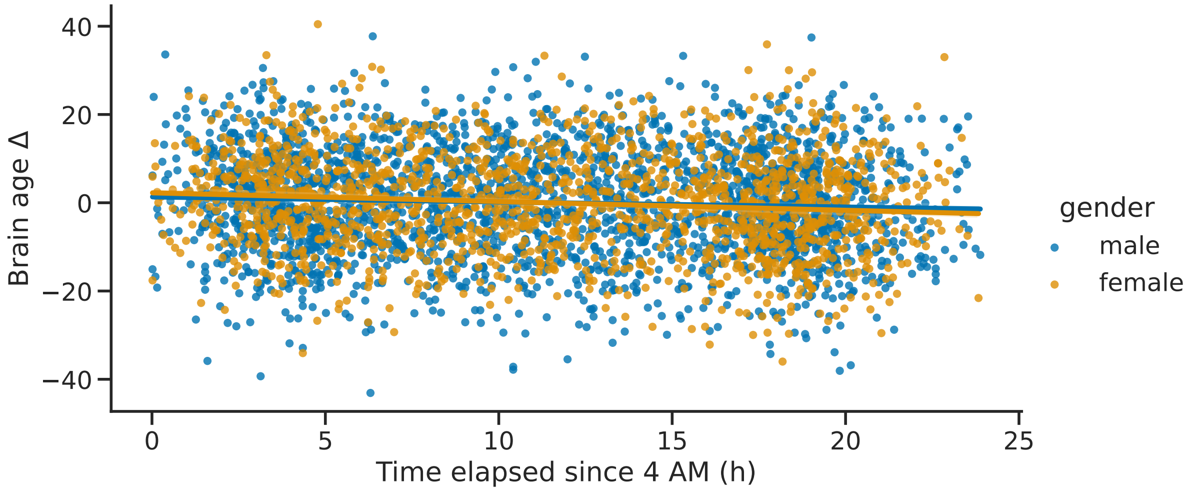


Figure S6: Effects of adjusted time of day (time elapsed since 4 AM) and gender on meditation-based brain age $\Delta$. The cross-validated age predictions obtained on MMD are shown as a function of the time at which recordings were started. Subject gender is color-coded (blue for male, yellow for female). Linear fits display the gender-specific impact of adjusted time of day on the brain age metric.

Alexandre Barachant, Stéphane Bonnet, Marco Congedo, and Christian Jutten. Classification of covariance matrices using a Riemannian-based kernel for BCI applications. *Neurocomputing*, 112:172–178, 2013. ISSN 0925-2312. doi: https://doi.org/10.1016/j.neucom.2012.12.039. URL `http://www.sciencedirect.com/science/article/pii/S0925231213001574`. Advances in artificial neural networks, machine learning, and computational intelligence.

Richard B Berry, Rita Brooks, Charlene E Gamaldo, Susan M Harding, Carole L Marcus, Bradley V Vaughn, et al. The AASM manual for the scoring of sleep and associated events. *Rules, Terminology and Technical Specifications, American Academy of Sleep Medicine*, 176, 2012.

Rajendra Bhatia, Tanvi Jain, and Yongdo Lim. On the Bures–Wasserstein distance between positive definite matrices. *Expositiones Mathematicae*, 2018.

Yilun Chen, Ami Wiesel, Yonina C Eldar, and Alfred O Hero. Shrinkage algorithms for MMSE covariance estimation. *IEEE Transactions on Signal Processing*, 58(10):5016–5029, 2010.

Alexandre Gramfort, Martin Luessi, Eric Larson, Denis A Engemann, Daniel Strohmeier, Christian Brodbeck, Lauri Parkkonen, and Matti S Hämäläinen. MNE software for processing MEG and EEG data. *NeuroImage*, 86:446–460, 2014.

Mainak Jas, Eric Larson, Denis A Engemann, Jaakko Leppäkangas, Samu Taulu, Matti Hämäläinen, and Alexandre Gramfort. A reproducible MEG/EEG group study with the MNE software: recommendations, quality assessments, and good practices. *Frontiers in neuroscience*, 12:530, 2018.

Fabian Pedregosa, Gaël Varoquaux, Alexandre Gramfort, Vincent Michel, Bertrand Thirion, Olivier Grisel, Mathieu Blondel, Peter Prettenhofer, Ron Weiss, Vincent Dubourg, Jake Vanderplas, Alexandre Passos, David Cournapeau, Matthieu Brucher, Matthieu Perrot, and Edouard Duchesnay. Scikit-learn: Machine learning in Python. *Journal of Machine Learning Research*, 12(Oct):2825–2830, 2011.

David Sabbagh, Pierre Ablin, Gaël Varoquaux, Alexandre Gramfort, and Denis A Engemann. Manifold-regression to predict from MEG/EEG brain signals without source modeling. In *Advances in Neural Information Processing Systems*, pages 7323–7334, 2019.

David Sabbagh, Pierre Ablin, Gaël Varoquaux, Alexandre Gramfort, and Denis A Engemann. Predictive regression modeling with MEG/EEG: from source power to signals and cognitive states. *NeuroImage*, page 116893, 2020.

J-B Schiratti, Jean-Eudes Le Douget, Michel Le van Quyen, Slim Essid, and Alexandre Gramfort. An ensemble learning approach to detect epileptic seizures from long intracranial EEG recordings. In *2018 IEEE International Conference on Acoustics, Speech and Signal Processing (ICASSP)*, pages 856–860. IEEE, 2018.
